# Modeling mosquito control strategies and their effect on pathogen transmission

**DOI:** 10.64898/2026.07.02.736114

**Authors:** Jacopo Rolfi, Andrea Radici, Claudio Bandi, Sara Epis, Paolo Gabrieli, Matteo Brilli

## Abstract

The mosquito Aedes *albopictus* is a competent vector for the transmission of several arboviruses and is currently spreading across many continents. Since conventional control methods, like insecticides, often lead to environmental problems and the emergence of resistance, scientists developed alternative mosquito control strategies. One of the most used is the Sterile Insect Technique (SIT), which involves the mass release of males sterilized through irradiation. The Toxic Male Technique (TMT) is instead based on the release of genetically modified males expressing toxic proteins that kill females when they mate. Control strategies are often intended as methods to eradicate mosquito populations, yet a less ambitious and more cost-effective task is to reduce them such that the probability of transmission of viruses to humans becomes negligible. To compare the efficacy of these control strategies, we develop a mathematical model with two communicating compartments: a mosquito population and epidemiological model coupled with a human epidemiological model. As a proof-of-concept, we test the model using meteorological and entomological data for the Emilia-Romagna region. Our results indicate that the TMT strategy is more effective in lowering the probability of transmission and provides indication for the deployment of control strategies.

## 1 Introduction

The tiger mosquito *Aedes albopictus* is an invasive species that was accidentally introduced in Italy from the United States in the late 1980s, likely through international trade routes [1, 2]. Among the mosquitoes present in Italy, *Aedes albopictus* is one of the most important vectors of diseases, carrying a wide range of pathogens, including viruses such as dengue, yellow fever, West Nile, Zika and chikungunya, and parasites such as *Plasmodium* spp. [3, 4].

Chikungunya virus (CHIKV) causes severe, debilitating and often chronic arthralgia, and was discovered in 1952 in Tanzania [5]. The history of CHIKV took a dramatic turn in 2004, when a new strain emerged in Kenya [6], spread to Indian Ocean islands, and later to Southeast Asia. During this period, thousands of infected travelers spread CHIKV into several regions of the world [7], resulting also in the first outbreaks in Italy [8], [9] and southern France [10]. These outbreaks shed light on the potential of CHIKV transmission in temperate regions, due to the ability of *Ae. albopictus* to survive in cold winters. Today, CHIKV infects millions of people throughout Europe, Asia, Africa and recently also Americas [11]. In 2025, 834 total cases of CHIKV were reported in France [12], and 469 in northern Italy [13], most of them in Emilia-Romagna [14].

Vector control measures are widely recognized as essential tools for reducing mosquito population density and, consequently, the transmission potential of pathogens [15]. When properly implemented, such interventions are among the most cost-effective public health measures; conversely, insufficient investment in vector control leaves the burden of vector-borne diseases unaffected [16]. Conventional, insecticide-based treatments are among the most used methods to control mosquitoes [17], despite the loss of effectiveness due to the evolution of resistance and the effects on non-target populations [18], raising concerns about the impact on environmental and human health [19].

Genetic biocontrol techniques (GBT) constitute the most promising alternative to insecticides. In this family of control methods, the sterile insect technique (SIT), invented by [20] to eradicate screw wormsin North America, is now actively used against mosquitoes. SIT relies on the release of a large quantity of males sterilized through X-ray irradiation. When females mate with these sterile males they produce eggs unable to hatch [21]. Despite high variation in hte way it has been applied, SIT has produced encouraging results in different regions of the world [22, 23]. A variation of this technique exploits the intracellular bacterium *Wolbachia* [24] a common intracellular symbiont of arthropods [25], which has been largely employed for vector control purposes, such as of *Ae. aegypti* [26]. *Wolbachia* causes an activation of the immunity response in insects [27], compromising the vectorial capacity of mosquitoes [28] and it leads to cytoplasmic incompatibility, resulting in reduced embryo survival and reduced hatching rates [29].

Others GBT have been proposed, such as the release of insect carrying a dominant lethal gene (RIDL) after engineering [30]. These transgenic insects, reared on a protective diet, are released in the environment where they mate with their wild relatives, transmitting the dominant lethal gene to the progeny that dies at the larval or pupal stage [31], [32].

Collectively, GBT strategies require at least one generation to affect the target population, since they do not directly affect the adult stage, but the offspring. This means that infected adult females can keep spreading pathogens also after the treatment.

The need of a timely biological control method led different works to look for approaches where the viability of the mated females could be affected [33, 34, 35, 36], where males are engineered to express toxic seminal fluid proteins (SFPs), These proteins, under natural conditions are produced in the male accessory glands (MAGs) and transferred to females during mating, reaching the female reproductive tract and haemolymph; also, they can interact with receptors in the female central nervous system [37, 38] and modulate post-mating responses, such as reduced receptivity to further mating and changes in female lifespan [39, 40, 41]. In this framework, males are engineered to deliver toxic SFPs that enhance these effects and negatively impact female survival. In 2025 Beach and Maselko [33], has been demonstrated that toxic SFPs do not affect male viability, contrary to what happens by irradiating males with X-rays in SIT. This latter trial, named by the authors Toxic Males Technique (TMT) can be defined as *intragenerational*, because it directly affects the life-span of adult females in the present generation thereby immediately reducing their ability to transmit pathogens.

Mathematical modeling plays a pivotal role in this context, since it provides a quantitative framework to assess the effectiveness of vector control strategies and the identification of key factors determining success or failure, prior to field implementation. In fact, mathematical models can simulate the dynamics of epidemic outbreaks in different environmental and weather conditions; indeed, over the past two decades, deterministic models based on ordinary differential equations (ODEs) have been increasingly used to simulate mosquito population dynamics and evaluate the effects of control strategies, beginning with SIT [42]. Moreover, the integration of vector population dynamics with models of pathogen transmission in humans, enables to focus on how different control strategies impact the spread of specific diseases in humans, beyond their direct effect on the abundance of mosquitoes.

In this study we adapt an age-structured mosquito population model developed by [43] to include SIT and TMT and we integrate it with a SEIR model to explicitly follow the propagation of infection from the mosquito to the human compartment and viceversa.

Instead of focusing on the mosquito population only, we consider how the control strategies affect epidemic spread in humans, with a first application on provinces of the Emilia-Romagna region, in Italy.

## 2 Materials and Methods

### 2.1 Mosquito population model

We derived the mosquito population model from [43]. The original model was validated with entomological data concerning Southern France and *Anopheles* species, but it was later applied to other geographical contexts and mosquito species, such as *Ae. albopictus* in Reunion Island [44, 45]. The ODE model (Eq. 1-7) takes as input daily temperature and rainfall levels to predict the abundance of multiple mosquito developmental stages; equations describe the development of eggs into larvae and pupae (E = eggs, L = larvae, P = pupae), which develop into emerging adult males (M) or females (F_*em*_). After an implicit blood meal, these latter become nulliparous females (F_*n*_). In turn, they develop into parous females (*F*_*p*_) after oviposition. We merged the original structure of the adult females classes (host-seeking, engorged and ovodepositon-site seeking) into nulliparous and parous females to avoid an over-parameterization of the model, as done in [44]. The presented mosquito system comprises Eq. 1-6 from [44], with minor modifications, while Eq. 7 was added to enable the implementation of the control strategies.

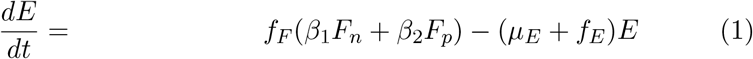

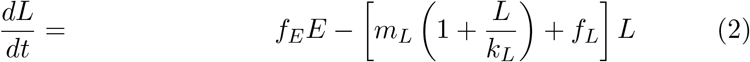

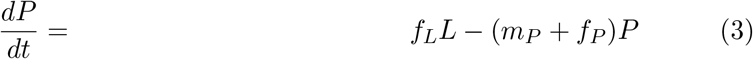

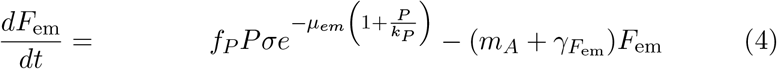

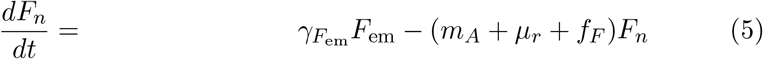

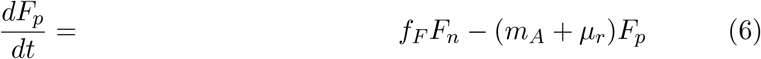

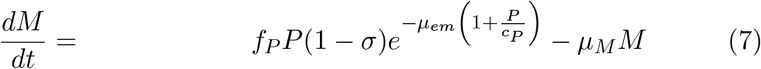

In the equations above, parameters are indicated by Greek letters, while functions (*d*^−1^) are in lower case Latin and depend on rainfall (*R*) and temperature (*T*).

Eq. 1 describes the dynamics in time of the abundance of eggs; *f*_*F*_ is the rate of transition from nulliparous to parous females, while *f*_*E*_ is the transition rate from eggs to larvae; these functions model the oviposition rate and are defined as:

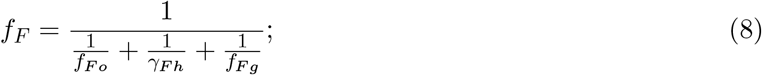

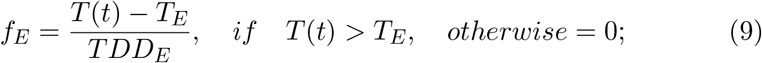

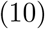

with:

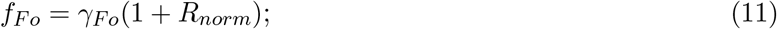

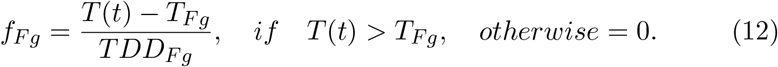

where R_*norm*_ represents the rainfall levels scaled to the range [0, 1]. The value of*f*_*E*_ at a certain time point is zero if the average temperature in the previous week was less than *T*_*E*_, and it linearly increases with temperature otherwise; at 25 °C *f*_*E*_ is 0.09, corresponding to an average duration of the stage of 1*/*0.09d^−1^ ≈ 11d. Instead, *f*_*F*_ combines transition rates (*f*_*F o*_, *γ*_*F h*_ and *f*_*F g*_) from implicit consecutive stages. The parameters of the model correspond to:

- *γ*_*Fo*_: development rate from ovipositing to host-seeking females;
- *γ*_*Fh*_: development rate from host-seeking to engorged females;
- *T*_*Fg*_: minimum temperature required for egg maturation;
- *T*_*E*_: minimum temperature for egg development;
- TDD_*Fg*_ total degree-days required for egg maturation;
- TDD_*E*_: total degree-days for egg development;
- *β*_1_: oviposition rate of nulliparous females;
- *β*_2_: oviposition rate of parous females;
- *μ*_*E*_: egg mortality rate.

and their values are reported in Supplementary Materials 1.

Eq. 2 describes the abundance of larvae, and depends on the following functions:

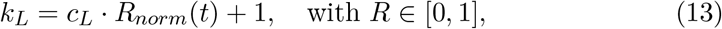

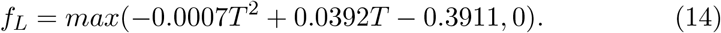

The *m*_*L*_ function governs the mortality of larvae and depends on temperature; it was obtained by modifying the corresponding mortality function proposed in [46] as 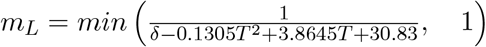. However, this formula assumes negative values outside the temperature range [− 6.54, 36.13] °C and has moreover a discontinuity when it changes sign. For this reason, we modified it as explained in Supplementary File 1. Along with the larval mortality, temperature affects the developmental rate to pupae and the corresponding function (*f*_*L*_) is zero outside the range [14.64, 41.36]°C.

Eq. 3 describes the dynamics of pupae, where *f*_*P*_ = *max*(0.0008*T* ^2^ − 0.0051*T* + 0.0319, 0) is the development rate from pupae to emerging adults. Similarly to the mortality rate for larvae (*m*_*L*_) function *m*_*P*_ was modified starting from the empirical formula 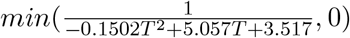, to avoid discontinuities and negative values outside the range [− 0.68, 34.35]°C (see Supp. File 1 for details). *k*_*P*_ is the environmental carrying capacity for pupae, is defined as:

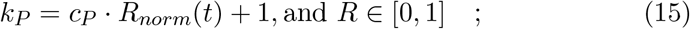

where *c*_*P*_ is a parameter estimated using field observations in Emilia-Romagna and published by [47]. We assume *c*_*L*_ = *c*_*P*_ as in [45].

Eq. 4 describes the dynamics for emerging adults females. The sex-ration *σ* is assumed to be 0.5. The intrastage competition, which affects emerging pupae [48], is accounted for by a pupal density-dependent term: 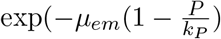; the mortality rate of adults is defined as:

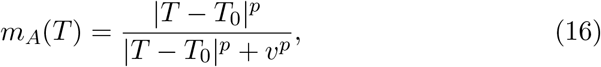

where at low and extremely high temperatures, *m*_*A*_ = 0.25, corresponding to a characteristic lifetime of ≈ 3 days, while at *T*_0_, *m*_*A*_ = 0.033, corresponding to ≈ 21 days; *p* and *v* were defined through calibration. The reference temperature *T*_0_ was fixed at 22°C based on experimental evidence reporting minimum adult mortality of *Aedes albopictus* at this temperature [49]. Consistently, the function attains its minimum at *T* = *T*_0_.

Finally, 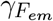 is the transition rate from emerging to nulliparous females. Eq. 5 models the abundance of nulliparous females, who have taken their first blood meal but that did not lay eggs yet, while parous females oviposited at least once (Eq. 6). In both equations, the mortality rate parameter is related to the seeking behavior (*μ*_*r*_).

To account for the dynamics of males and enabling the implementation of the control strategies, eq. 7 was added in this work,. Dynamics of males are similar to that of emerging females, but include only mortality (*μ*_*M*_).

Rainfall plays a key role in the model, especially for aquatic stages (eggs, larvae and pupae). Indeed, the presence of water has the effect of triggering the hatching of eggs [50], but heavy rains can also flush the available breeding sites [51], influencing the mortality rate of larvae and pupae.

Based on the above model and the available literature [44], we defined the mathematical implementation of the control strategies (See Supplementary File 1). For both SIT and TMT - indicated by subscript *s* for sterile and *k* for toxic (to avoid using letter *t*) -, simulations include *n* releases of *λ* males every *τ* days, implemented in the model as time-dependent events.

According to recent research, lab-reared males have increased mortality rate (*μ*_*s*_) compared to their wild counterpart [52, 53] and reduced ability to find/attract/mate with females [54]. Therefore, we modulated the probability for an emerging female to mate with a released male as:

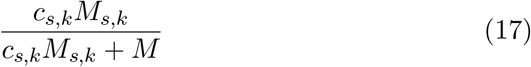

where *c* is the reduced competitiveness *c*_*s,k*_ ∈ [0, 1] of lab-reared males (*c*_*wild*_ = 1). Recent studies highlighted that sterilization of males with the sex pheromone heptacosane can instead have *c > c*_*wild*_ [55]. While we run all simulations with *c* = 0.5, our code enable to easily test alternative scenarios. In the case of TMT, there is no information about reduced male competitiveness, but we kept a reduced competitiveness as for sterile males to avoid introducing a bias in the analysis.

### 2.2 Integration of the mosquito and the human compartment models

In this section, we describe the integration of a SEIR model into the previously described mosquito population model (Figure 1). The interaction of the mosquito and human compartment is provided by pathogen transmission from human to mosquitoes and viceversa via biting. In equations, we denote human compartments by subscript *h*; *S*_*h*_, *E*_*h*_, *I*_*h*_ and *R*_*h*_ are therefore the abundance of the susceptible, exposed, infected and recovered compartment, respectively. We partition adult females (*F*_*n*_ and *F*_*p*_) into pathogen-free, exposed, and infectious; the last two classes are denoted by subscripts *e* and *i*, respectively. The exposed class in both the mosquito and human compartments captures the incubation period (extrinsic, or EIP, for mosquitoes, indicated by parameter *η*; and intrinsic, or IIP, for humans, indicated by *ϵ*), during which the pathogen develops within a non-infectious mosquito or human [56]. We assumed negligible mortality for humans, whose total abundance remains constant (*N*_*h*_(*t*) = *N*_*h*_). therefore we removed the equation for *R*_*h*_ since it can be obtained from the other variables (*R*_*h*_(*t*) = *N*_*h*_ − (*S*_*h*_(*t*) + *E*_*h*_(*t*) + *I*_*h*_(*t*))). The overall model equations are:

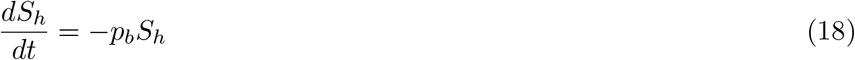

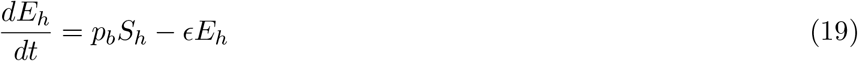

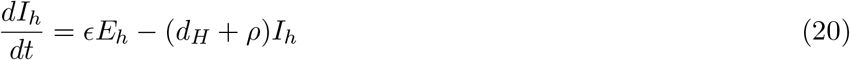

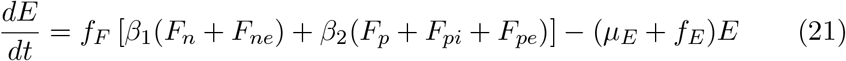

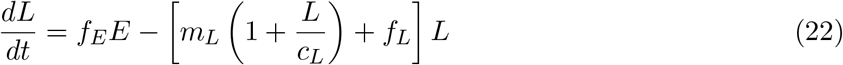

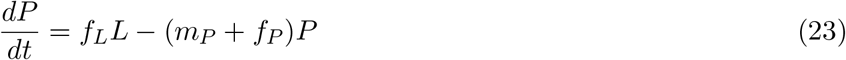

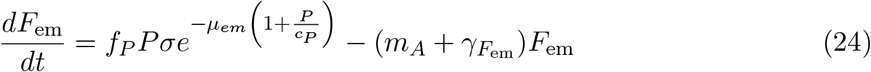

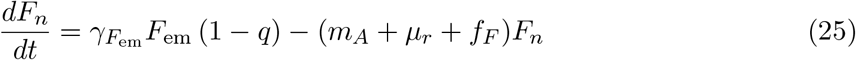

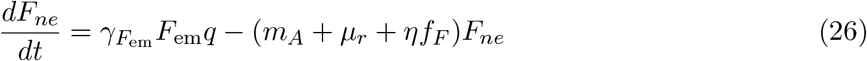

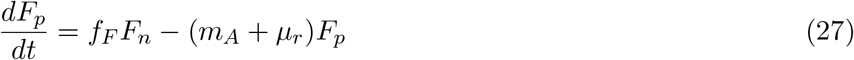

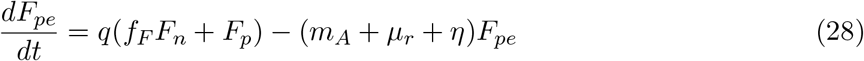

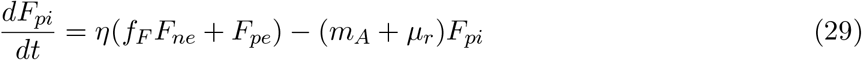

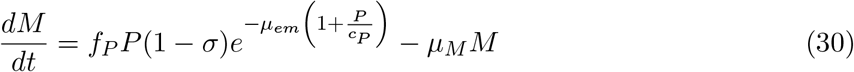

**Figure 1.**
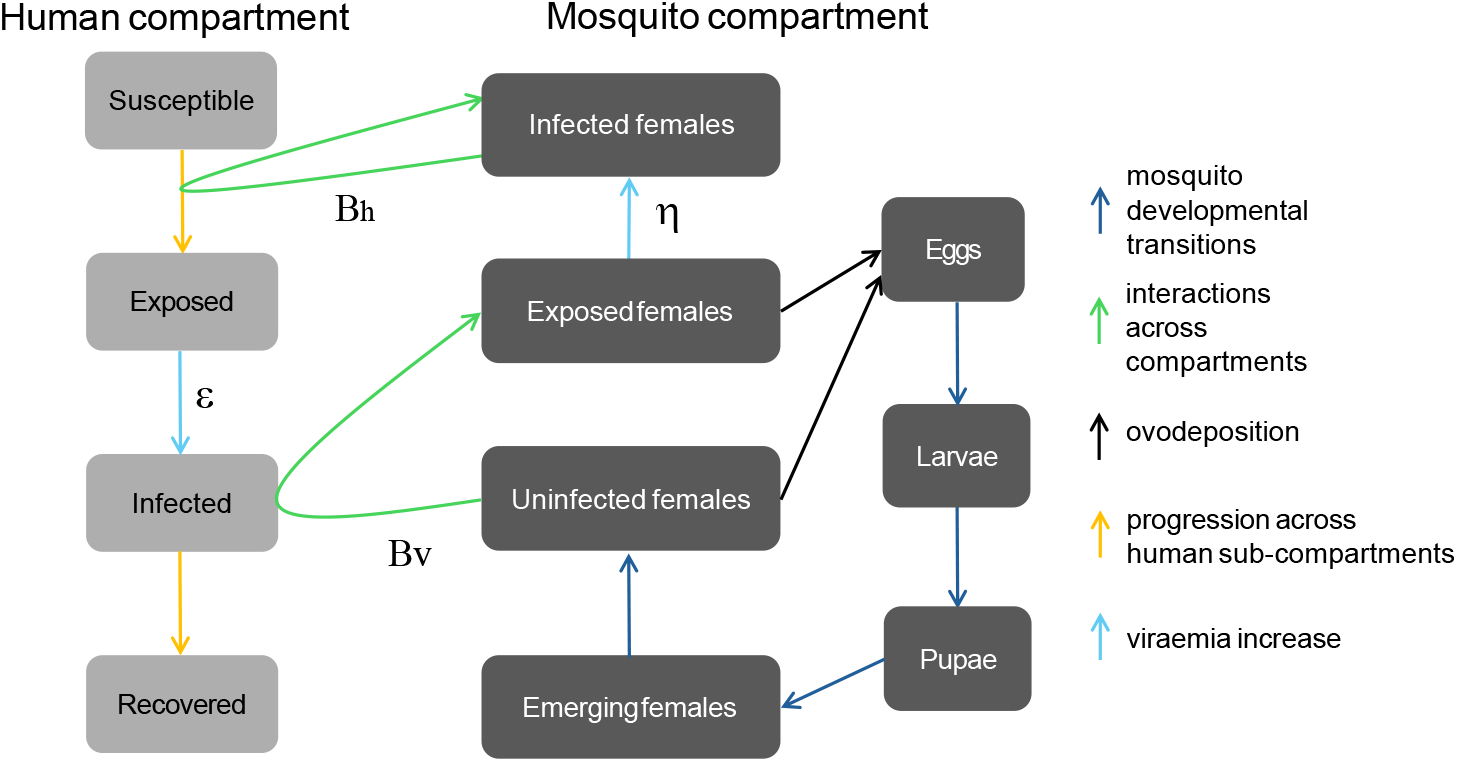
Simplified scheme of the model showing the Human and Mosquito compartments and their interactions.

The transition from susceptible to exposed humans depends on the probability of being successfully bitten by an infectious mosquito (*p*_*b*_), defined as:

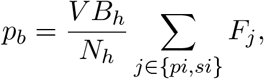

where *V* is the mosquito biting rate and *B*_*h*_ is the transmission probability of the pathogen when a mosquito bites a human. The sum at the numerator runs over all possible infected classes of mosquitoes.

The probability for an adult female mosquito to become infected after biting an infected human is given by:

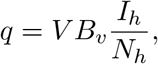

Where *B*_*v*_, the transmission rate of the virus from human to mosquito. *B*_*v*_, *B*_*h*_ and *V* are Brière functions [57]:

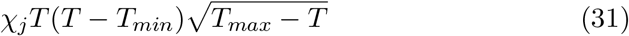

where *χ*_*j*_ is a constant controlling the maximum of the function, *T*_*min*_ is the minimum and *T*_*max*_ maximum critical temperatures, respectively. The value of the function is forced to zero for *T < T*_*min*_ and *T > T*_*max*_ (See Supplementary File 1).

### 2.3 Male release strategies

To implement a sounding mosquito control scheme, we need to define the starting date of releases (*t*_*start*_), their total number (*n*), the interval between two releases (*τ*) and the number of males per release. When designing field vector control operations, these quantities are defined on the basis of a compromise between effectiveness and economical and technical constraints. Releases are usually limited by the total allocated budget and the technical limits of mosquito rearing in labs. In this study, we defined a parameter space for *t*_*start*_, *n* and *τ* by assigning a plausible range to each of those parameters, and then we explored the consequent dynamics predicted by the model. In total, considering both control strategies, we simulated 108 scenarios (Figure 4). For each scenario, we computed the corresponding average basic reproduction number *R*_0_, as defined in Eq. (32). This way, we identified the combination of parameters leading to the strongest reduction in *R*_0_(*t*) with respect to the standard (no treatment) scenario.

Specifically, the simulations were performed by progressively delaying the first release date (*t*_*start*_), by beginning in April up to July. This parameter was increased by 15 days in each simulation. This temporal window was selected because in temperate regions *Ae. albopictus* populations typically reach their highest abundance during the spring–summer period. Within this window, we defined both the total number of releases (*n*) and the time interval between consecutive releases (*τ*), based on realistic operational constraints and the deployment capacity of the facilities responsible for implementing the interventions [22]. Furthermore, the number of males released at each impulse (*λ*) was determined based on the mean simulated annual peak of female abundance using the standard model.

Here, the competitiveness of sterile/toxic males parameter can be considered as a scaling factor of the total abundance of released males *λ* (Eq. 17), therefore, when testing different values of *λ*, we are implicitly testing different competitiveness values. For instance, *λ* can be interpreted as an effective number of sterile/toxic males whereas the actual number of released males would be *λ/c*: releasing 5 × 10^5^ males with *c* = 1 corresponds to the release of 1 × 10^6^ males with *c* = 0.5 or 2 × 10^6^ males with *c* = 0.25. Across our simulations, we always used *c* = 0.5.

#### 2.3.1 Insecticide treatment

In Section 3.3, we studied a situation where control strategies are deployed after the detection of an infected individual (*patient 0*), with some delay. In recent vector control operations in Italy, this often meant that a certain area around *patient 0* gets treated with adulticides. These localized treatments strongly reduce the abundance of mosquitoes in a relatively small area (≈ 200 × 200*m*^2^) but barely affect those nearby. Following treatment, mosquitoes from surrounding areas rapidly recolonize the treated zone, causing population abundance to return to pre-treatment levels within a few days. Our model does not explicitly account for spatial connectivity of mosquitoes; therefore, modelling insecticide applications only as an increase in mosquito mortality would lead to a substantial overestimation of the impact of treatments. We therefore model the repopulation effect indirectly, interpreting the local and transient reduction in mosquito abundance as a temporary decrease in the probability that a mosquito acquires infection when biting an infectious host. Consequently, in our model we implement the following scheme: *i)* at a given date, *patient 0* is detected; *ii)* with some delay, we remove 0.99 × *S* × *A* mosquitoes, with *S* being the proportion of the treated area (*S* = 0.2) and *A* all adult classes in the mosquito model; *iii)* concomitantly, we multiply *q*, the probability that mosquitoes get the infection, by 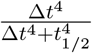, where Δ*t* is the time elapsed from treatment and *t*_1*/*2_ = 7 days is the Δ*t* such that *q*_*treat*_ = *q/*2 (Figure 2).

**Figure 2.**
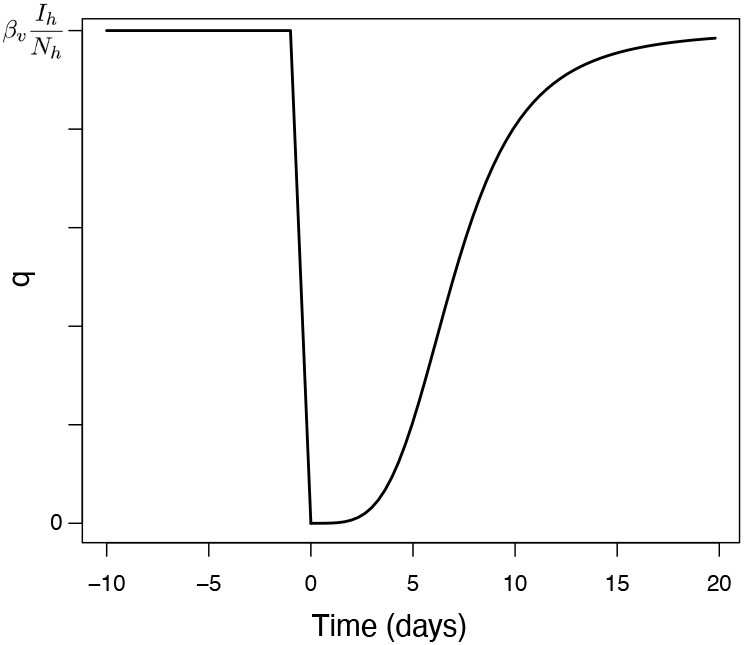
Illustration of the implemented reduction in *q* after an insecticide treatment at *t* = 0.

### 2.4 Epidemiological indicators

In this study, we aim to compare the effect of different control strategies on the ability of female mosquitoes to transmit a virus to the human compartment; consequently, we focused on the following indicators:

- The basic reproduction number 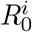, computed for each scenario *i* ∈ *s, k, c* (sterile, killer, control), based on [58, 59]:

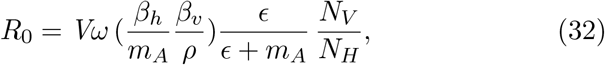

that incorporates terms related to: the biting rate; human host preference; the average number of infected humans by a single infective mosquito; the average number of mosquitoes infected by the introduction of a single infective human into the population; the mosquito mortality function; human recovery rate; mosquito extrinsic incubation period of the virus; total number of humans; total number of adult females mosquitoes; (see Supplementary File 1).
- The *relative reduction* of the basic reproduction number with respect to the standard scenario, defined as:

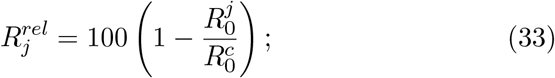

Mosquito populations is initalised to a value of 100 eggs at the beginning of each simulation (*t = 0*), with a spin-up period of 3 years. Furthermore, we calibrated the model by minimizing the discrepancy between simulated and observed egg counts using a bounded nonlinear optimization procedure (the `nlminb` function in R). The objective function minimizes the root mean squared error computed on normalized egg-count time series, varying a subset of 24 parameters to which the model is most sensitive (out of 46 total parameters).

### 2.5 Meteorological and entomological data

In this study we focused on the Emilia-Romagna provinces of Forlì-Cesena, Rimini, and Ravenna (Figure 3), a biogeographical area classified as continental [60]. We computed the daily mean temperature for each of the considered provinces by downloading the Arpae Emilia-Romagna data from the `dext3r` open sources portal [61], using all the available weather stations in the study area; rainfall data (mm) were retrieved from E-OBS gridded dataset (at 0.1° × 0.1°; Figure 3). To avoid introducing discontinuities in the data during integration and to have data for all integration points, we input meteorological data to the model using the R function `approxfun`.

**Figure 3.**
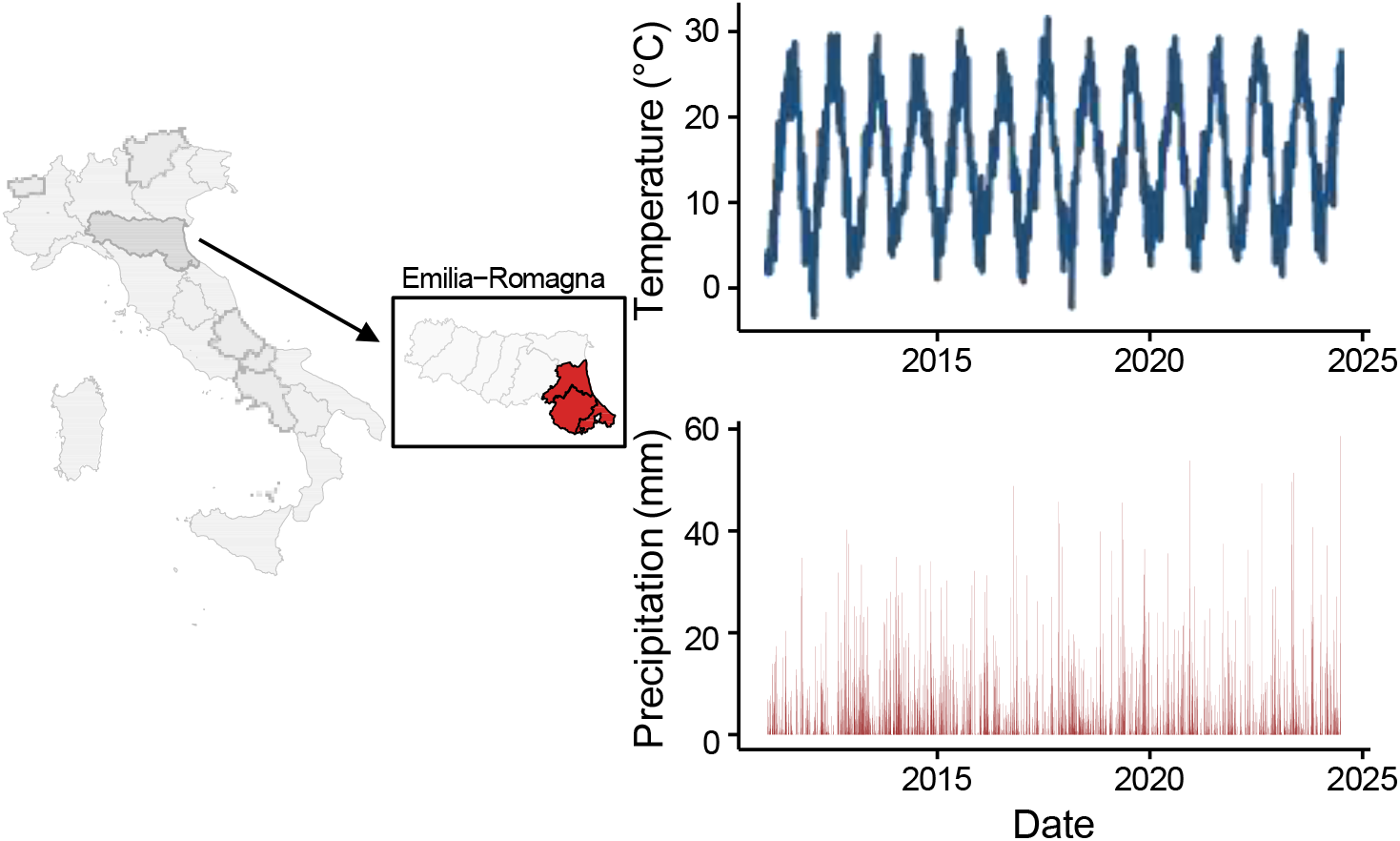
Region of study and corresponding meteorological data for the entire period.

We used ovitrap data available on VectAbundance v0.15 database [62] for the years 2010 − 2022 and the Emilia-Romagna region [63, 64]. Each observation in this dataset represents the cumulative week sum of eggs per ovitrap. The dataset accounts for microclimatic variation and sampling differences, by aggregating the data to a 9 × 9 km grid using the median ovitrap value within each grid cell at each observation date (for further details, see [62]). We used the RStudio 2025.9.0.387 version (http://www.rproject.org/) for all computations.

## 3 Results

### 3.1 Definition of parameters for treatment

By extracting 54 triplets from the intervention parameter space, we defined our intervention strategies for both control strategies. In this phase, we fixed the number of released males at a number similar to the number of adult female mosquitoes observed at the peak i.e. 1.5 × 10^5^.

In Figure 4 we show the average *R*_0_ relative to CHIKV transmission for each tested scenario, for each control strategies, averaged over 9 years. The 9-years average allows to smooth the large interannual fluctuations due to different temperature and rainfall levels affecting the abundance of mosquitoes and compare control strategies in a robust way (Table 1).

**Table 1.** Best scenarios identified for the two strategies.

| Method | $t_{\text{start}}$ | $n$ | $\tau$ | $R_0$ |
| --- | --- | --- | --- | --- |
| SIT | yyyy-05-13 | 15 | 5 | $5.24 \pm 1.68$ |
| TMT | yyyy-05-28 | 15 | 5 | $3.45 \pm 1.18$ |

**Figure 4.**
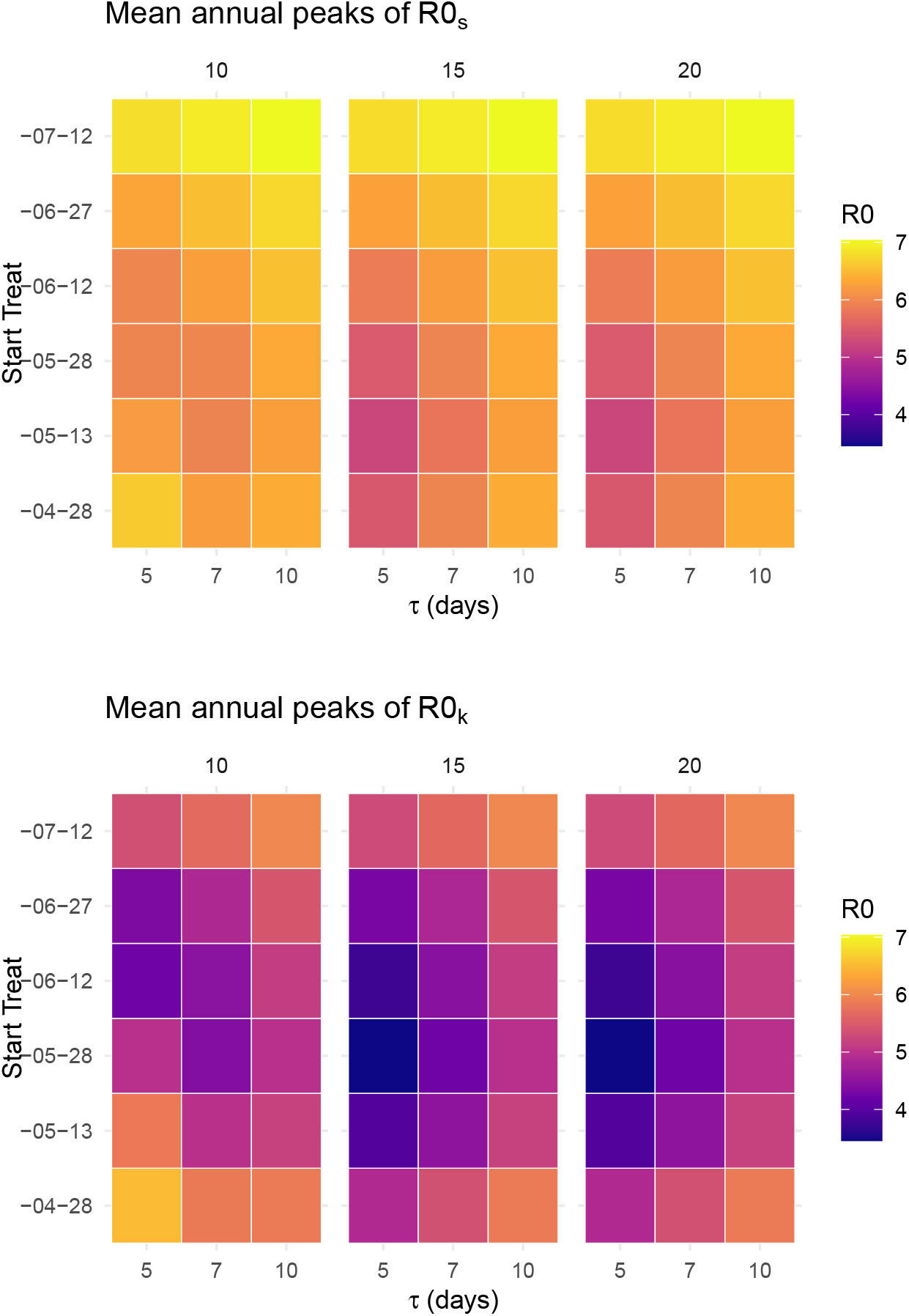
Mean R_0_ annual peaks. Upper panel, SIT; lower panel, TMT. *τ* is the interval separating releases, *Start Treat* defines the date at which treatment starts. Each of the six panels represent the total number of releases during the year *n*.

### 3.2 Scenario 1. Control strategies for prevention

The analysis of the *R*_0_ time series (Figure 5a) highlights a reduction induced by both control strategies with respect to the standard scenario. The maximum *R*_0_ remains above the theoretical epidemic threshold (*R*_0_ = 1) in all cases (Figure 5b).

**Figure 5.**
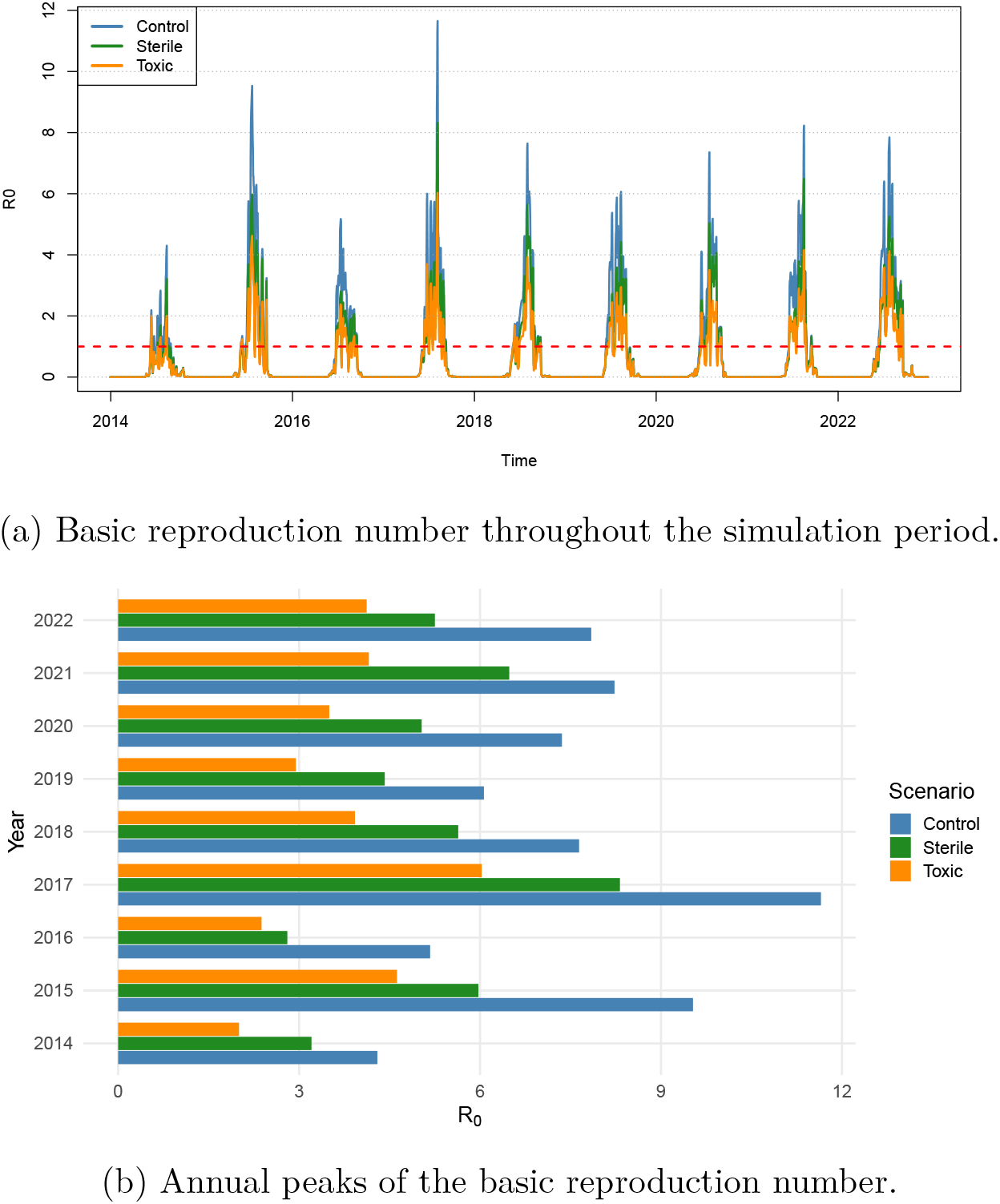
Temporal evolution and annual peaks of the basic reproduction number under different control scenarios.

To test if there is any chance of reducing *R*_0_ below the epidemic threshold, we simulated a gradual increase of the value of *λ*. To better illustrate the differences of the two techniques relative to the standard scenario, we calculated the percentage of reduction of the basic reproduction number over the entire time span for the above simulations (Eq. 33). In agreement with the previous results, TMT always results in a stronger reduction of the basic reproduction number for all pulse sizes (Figure 6, averages throughout years in Tables 2, a and b). Since the average *R*_0_ for the standard scenario is 7.53, reductions over ≈ 86% correspond to basic reproduction numbers below the epidemic threshold; our analysis shows that TMT is systematically closer to the epidemic threshold condition than SIT for each value of *λ* tested, but only in two years and at the maximum tested *λ*.

**Table 2.**
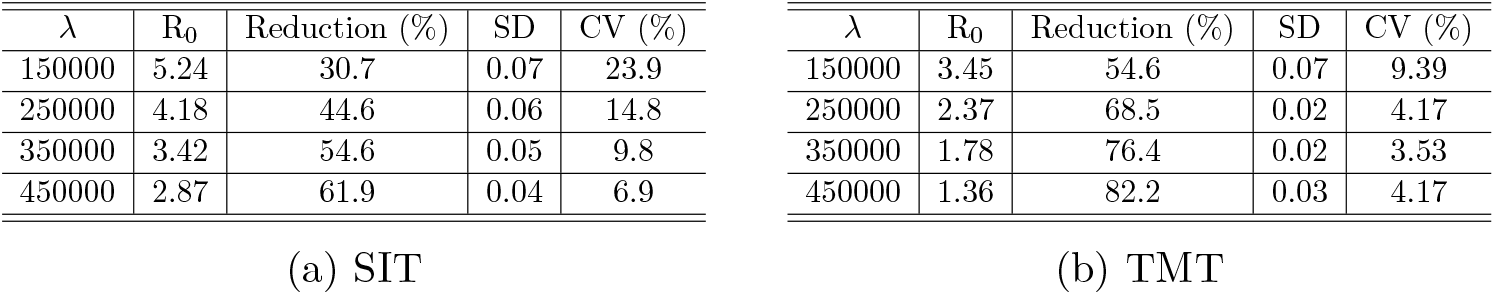
Mean reduction in *R*_0_ across the study period under the two control strategies. For each value of *λ*, the mean reduction rate was first computed for each year; standard deviation (SD) and coefficient of variation (CV) were then calculated over all years.

**Figure 6.**
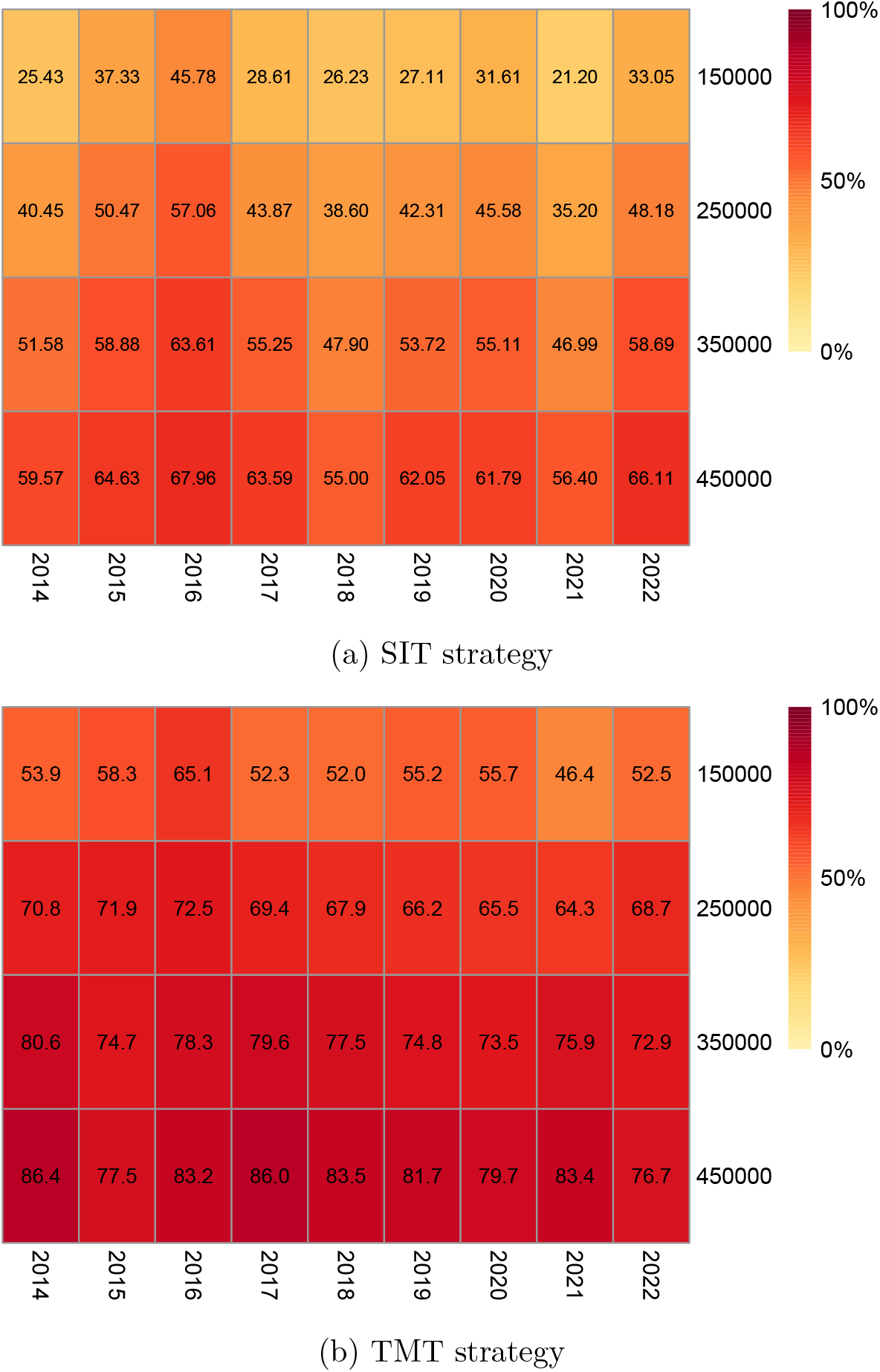
Heatmaps of the reduction in the peak basic reproduction number as a function of the number of males released per pulse (*λ*). Rows correspond to different values of *λ*, while columns represent intervention years.

Finally, we wanted to assess the difference in the epidemiological dynamics of the infection in the human compartment, in the standard situation and the two control strategies, using a fixed value of *λ* = 1.5 × 10^5^. The simulations were performed by initializing the human compartment with *I*_*h*_(*t*_*i*_) infected individuals and *S*_*h*_*t*_*i*_ = *N*_*h*_ − *I*_*h*_(*t*_*i*_) for a given date *t*_*i*_. Infected individual(s) is(are) introduced a month after the first release in the case of simulations of the control strategies; for the standard scenario, the infected indivdual(s) is (are) introduced on June 15^*st*^. This approach corresponds to the arrival of one or more infected individuals in the study area, for instance after a having traveled to a region where CHIKV is endemic. At the beginning of the simulation, all mosquitoes are susceptible. The ensuing dynamics are summarized in Figure 7 and in Table 3.

**Table 3.** Fraction of humans getting infected throughout the entire simulation.

| Initial $I_h$ | $\sum I_h$ (%) | Initial $I_h$ | $\sum I_h$ (%) | Initial $I_h$ | $\sum I_h$ (%) |
| --- | --- | --- | --- | --- | --- |
| 1 | 99.2 | 1 | 35.6 | 1 | 10.8 |
| 5 | 99.8 | 5 | 79.1 | 5 | 39.7 |
| 10 | 99.9 | 10 | 90.3 | 10 | 58.8 |
| 15 | 99.9 | 15 | 93.9 | 15 | 69.5 |
| 20 | 99.9 | 20 | 95.6 | 20 | 76.1 |

(a) Standard (b) SIT (c) TMT

**Figure 7.**
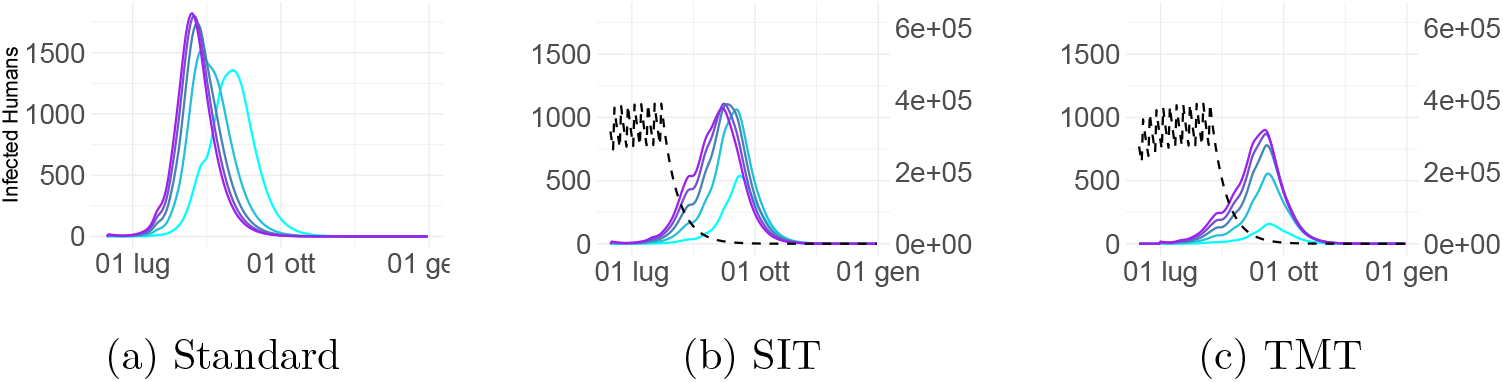
Dynamics of infection in the human compartment. Left axis: infected humans, continuous lines; right axis in (b) and (c): released males, dashed line. Colours of series represent initial infection seeding levels (1, 5, 10, 15, 20), mapped onto a cyan-to-purple gradient from low to high initial infection.

As expected, the infection dynamics are highly sensitive to the initial number of infected humans (*I*_*h*_) (Figure 7). Only the TMT scenario enables a relatively strong reduction of the number of infected humans, whereas SIT loses very rapidly its efficiency as the number of initial infected increases (Table 3).

### 3.3 Scenario 2. Post-detection control

In reality, insecticide? treatments are more often applied only some days after infected individuals are detected, for instance when they start presenting symptoms and get in touch with the family physician or get hospitalized. Therefore, we simulated the dynamics of the infection by introducing a delay in the treatments (3, 7 or 10 days) after the first infected human enters the population. In this scenario, we did not account for the emergence of resistant strains, which is instead pretty common when treatments with insecticides are repeated in time [65]. Here, the infected human enters the population at the peak of the mosquito population, and treatment occurs with the indicated delay (Figure 8).

**Figure 8.**
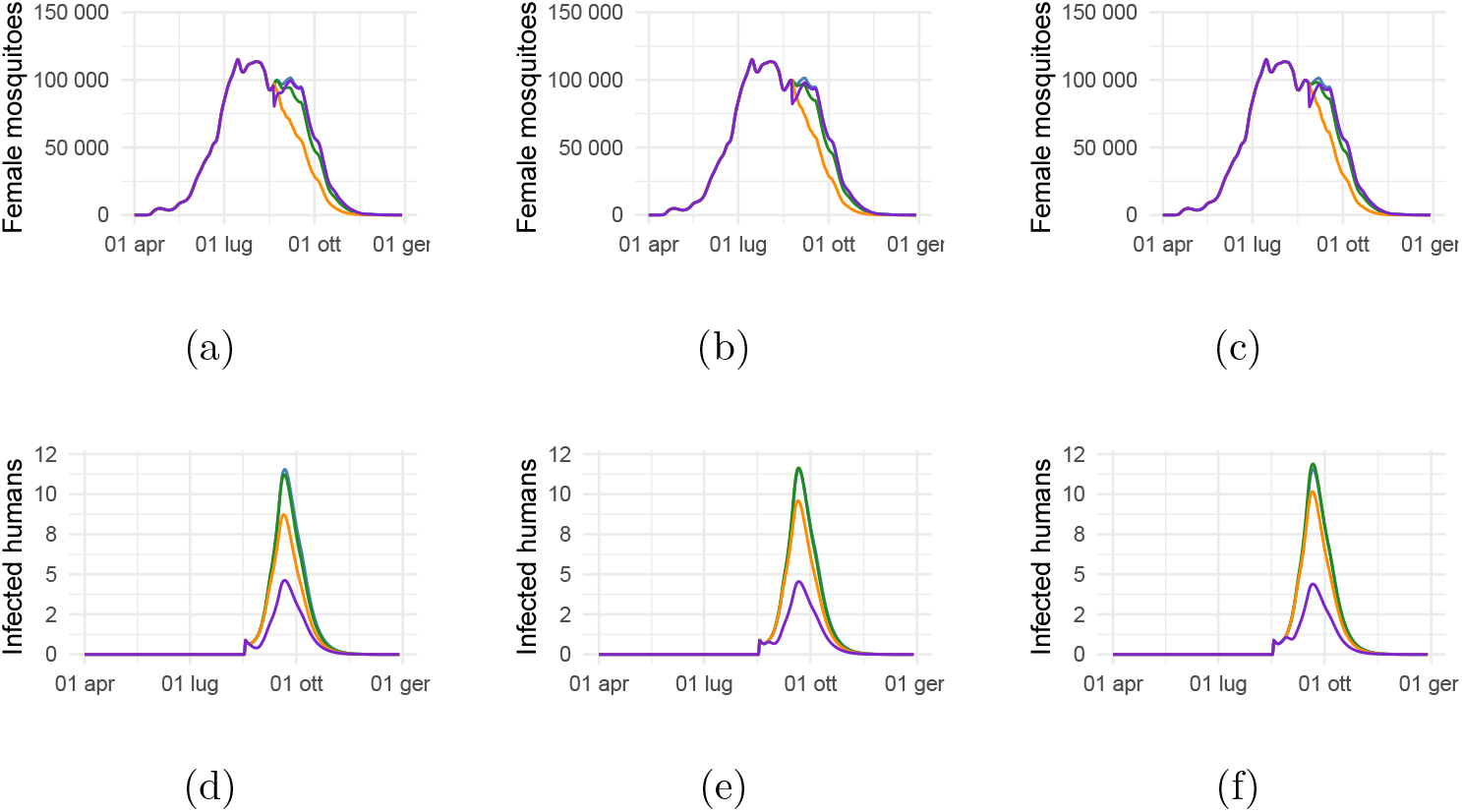
Adult females mosquitoes (a, b, c), and corresponding infected humans (d, e, f). Delay in treatment: first column 3, second column 7, third column 10 days. Colors indicate the type of treatment: blue, standard situation; purple, insecticide; green, SIT; orange, TMT.

Our simulations show that both the dynamics of adult female mosquitoes and infectious humans are more significantly influenced by the type of treatment, rather than the delay. In this analysis, TMT has similiar effects of SIT considering humans and mosquitoes compartments, instead insecticide treatment outperforms both strategies considering the virus dynamics in humans compartment.

## 4 Discussion and conclusion

In this work, we compared vector-control strategies by their ability to reduce epidemic permissiveness and to mitigate outbreak size following viral introduction, rather than by their ability to decrease mosquito abundance *per se*. We started with a published weather-driven mathematical model of mosquitoes [44], we adapted it to make it usable with environmental data specific of the geographical area under investigation and then we implemented two mosquito control strategies, namely Toxic Male [33] and Sterile Insect technique [66, 67, 68]. Adaptations to the geographical area were necessary because several temperature-dependent functions did not behave meaningfully with our temperature range, for instance by changing sign or breaking down. Next, we coupled the mosquito model with a SEIR model to follow the propagation of a viral disease, specifically CHIKV in both the human and *Aedes* compartments. The final aims of this study were *(i)* to obtain an adaptable tool to support the decision-making process of public health authorities when planning mosquito control strategies, *(ii)* to quantify the effectiveness of SIT and TMT relative to a no-control scenario and *(iii)* to explicitly follow the epidemiological dynamics of humans and mosquitoes.

Our results indicate that TMT is more effective in both controlling mosquito abundance and viral spread across a wide range of intervention scenarios. In our analysis, *R*_0_ is always smaller under the TMT scenario suggesting that to have comparable outcomes with SIT one has to release more males, which is necessarily associated with increased costs, under the assumption that sterile and toxic males have the same cost. We found that the timing of releases is a key determinant of control efficacy. Because TMT acts directly on the adult female population, interventions can be initiated later in the season than SIT and other conventional strategies, for which earlier deployment is generally required [69].

Based on *R*_0_, we determined that the best starting date for both control strategies is before the annual mosquito abundance peak, when few wild competitors are present, such that the probability that released males mate with wild females is increased, in agreement with results by Haramboure et al. [44]. The outcome is strongly influenced by two key parameters related to treatments: the number of males released per intervention (*λ*) and their competitiveness (*c*). The former impacts on the cost of treatments, and cannot therefore be increased at will; even when cost is not a problem, there usually are technical limits in production rates. Competitiveness is instead a limit related to the rearing of males in artificial conditions, and can be likely underestimated in our simulations, especially for TMT. In previous works Haramboure et al. [44] several parameters related to the mosquito compartment, such as adult mortality, egg hatching rate ecc. were demonstrated to be instrumental for the success of control programs, for this reason we did not explore their role in our simulations.

In contrast to the results reported by Beach and Maselko [33], none of the simulated scenarios in our study led to mosquito population eradication. This difference can be attributed to both biological and modelling factors. In particular, the release ratios and environmental conditions considered here were insufficient to drive the population to extinction. The aim of our study was, however, to determine feasible control strategies capable of reducing arbovirus transmission risk using operational parameters (*λ, τ*, *n*) that might be realistically implemented by local facilities, without overestimating their deployment capacity. Moreover, since our model is formulated as a deterministic continuous-time system, population densities may approach zero asymptotically without reaching it exactly; this result indicates that both strategies, with this configuration of parameters, are likely unable to impeed the virus transmission to the human compartment.

When modeling insecticide treatment, we highlight that they are better than any other control strategy. A limitation is however the necessarily simplified representation of this highly local treatment; relevant factors are not explicitly considered, such as the spatial allocation of treatments, where smaller spatial scales models can account for the mobility of mosquitoes, and thus, a possible recolonization of the treated area can be modeled with more accuracy [70]. Furthermore, insecticides result in the selection of resistant genotypes, especially among mosquitoes [71], which reduce the efficacy of next treatments. In parallel, insecticides have side effects on non-target species, and are also responsible for other environmental and human-health concerns [18]. Therefore, even if our simulations show that insecticides application outperforms the other control strategies, in line with other works [72] we note that the above side-effects could counterbalance the benefits.

Recently, SIT and TMT have been proposed as preventive strategies rather than emergency control measures. By implementing them prior to the detection of a disease, they could help keeping the mosquito population low and significantly decrease the probability that a disease spreads in the human compartment, and this is clear from our simulations even if neither control strategy seem to extend its effect long after the last release. In contrast, our simulations show that when SIT and TMT are applied after the detection of infected individuals, they have a limited protective role, especially for SIT.

Ae. albopictus populations are strongly regulated by environmental variables, especially in temperate regions [73]. A promising direction for future research is the development of adaptive forecasting frameworks in which mechanistic population models are continuously updated using real-time meteorological observations and weather predictions. This would enable dynamic optimization of vector control strategies based on anticipated mosquito abundance and seasonal conditions.

These findings reinforce the value of mathematical models, capable of simulating the timing and magnitude of mosquito population peaks. Such tools could improve the scheduling of control interventions by providing short-term (1 − 4 week) forecasts of population dynamics, as demonstrated by Ventura et al. [74].

## Acknowledgements

JR and MB want to thank Luis Almeida (Sorbonne Univerisity) and Tiziano Penati (Università degli Studi di Milano) for helpful discussion on the paper and model. JR wish to thank project GSA-IDEA of the University of Milan for financial support. Partial support by NextGeneration EU-MUR PNRR Extended Partnership initiative on Emerging Infectious Diseases (Project no. PE00000007, INF-ACT) and Multilayered Urban Sustainability Action (Project no. ECS00000037 - MUSA).

