## Supplementary File 1 for "Modeling mosquito control strategies and their effect on pathogen transmission"

### 1 Models

#### 1.1 Sterile-males model

Here, we define the mathematical framework of the SIT, where M$_s$ stands for sterile males.

$$\begin{cases} \dfrac{dE}{dt} = f_F(\beta_1 F_n + \beta_2 F_p) - (\mu_E + f_E)E \\ \dfrac{dL}{dt} = f_E E - \left[m_L\left(1 + \dfrac{L}{c_L}\right) + f_L\right] L \\ \dfrac{dP}{dt} = f_L L - (m_P + f_P)P \\ \dfrac{dF_{\text{em}}}{dt} = f_P P \sigma e^{-\mu_{em}\left(1+\frac{P}{c_P}\right)} - (m_A + \gamma_{F_{\text{em}}})F_{\text{em}} \\ \dfrac{dF_n}{dt} = \gamma_{F_{\text{em}}} F_{\text{em}} \left(1 - \dfrac{cM_s}{cM_s + M}\right) - (m_A + \mu_r + f_F)F_n \\ \dfrac{dF_s}{dt} = \gamma_{F_{\text{em}}} F_{\text{em}} \dfrac{cM_s}{cM_s + M} - (m_A + \mu_r + f_F)F_s \\ \dfrac{dF_p}{dt} = f_F F_n - (m_A + \mu_r)F_p \\ \dfrac{dM}{dt} = f_P P(1-\sigma) e^{-\mu_{em}\left(1+\frac{P}{c_P}\right)} - \mu_M M \\ \dfrac{dM_s}{dt} = -\mu_{M_s} M_s, \end{cases} \tag{1}$$

#### 1.2 Toxic males model

Here we present the mathematical framework which models the dynamics of the population when accounting for the gene-drive technique:

$$
\begin{cases}
\dfrac{dE}{dt} = f_F(\beta_1 F_n + \beta_2 F_p) - (\mu_E + f_E)E \\
\dfrac{dL}{dt} = f_E E - \left[m_L\left(1 + \dfrac{L}{c_L}\right) + f_L\right] L \\
\dfrac{dP}{dt} = f_L L - (m_P + f_P)P \\
\dfrac{dF_{\text{em}}}{dt} = f_P P \sigma e^{-\mu_{em}\left(1+\frac{P}{c_P}\right)} - (m_A + \gamma_{F_{\text{em}}})F_{\text{em}} \\
\dfrac{dF_n}{dt} = \gamma_{F_{\text{em}}} F_{\text{em}} \left(1 - \dfrac{cM_k}{cM_k + M}\right) - (m_A + \mu_r + f_F)F_n \\
\dfrac{dF_p}{dt} = f_F F_n - (m_A + \mu_r)F_p \\
\dfrac{dM}{dt} = f_P P(1-\sigma)e^{-\mu_{em}\left(1+\frac{P}{c_P}\right)} - \mu_M M \\
\dfrac{dM_k}{dt} = -\mu_{M_k} M_k,
\end{cases}
\tag{2}
$$

##### 1.3 Parameters and functions

In this chapter, we present the parameters (in Greek) and the functions (in Latin) used for this study.

###### 1.3.1 Tables of parameters

Table 1: Parameters and functions of the mosquitoes-populations models.

| Notation | Description | Expression | Reference |
|---|---|---|---|
| $\beta_1$ | Number of eggs laid per ovipositing nulliparous female | 60 | [1] |
| $\beta_2$ | Number of eggs laid per ovipositing parous female | 80 | [1] |
| $\sigma$ | Sex ratio at emergence | 0.5 | Optimization |
| $\mu_e$ | Minimum egg mortality rate (day$^{-1}$) | 0.01 | Optimization |
| $\mu_{em}$ | Mortality rate during emergence (day$^{-1}$) | 0.1 | [2] |
| $\mu_r$ | Mortality rate related to host-seeking behaviour (day$^{-1}$) | 0.8 | Optimization |
| $\mu_M$ | Mortality rate of wild males (day$^{-1}$) | 0.0735 | [3] |

Continued on next page

| Notation | Description | Expression | Reference |
|---|---|---|---|
| $T_E$ | Minimum temperature required for egg development (°C) | 15 | Optimization |
| $TDD_E$ | Total degree-days required for egg development (°C) | 110 | [2] |
| $\gamma_{Fem}$ | Development rate of emerging females (day$^{-1}$) | 0.4 | [2] |
| $\gamma_{Fo}$ | Transition rate from ovipositing to host-seeking females (day$^{-1}$) | 0.12 | Optimization |
| $\gamma_{Fh}$ | Transition rate from host-seeking to engorged females (day$^{-1}$) | 0.3 | Optimization |
| $T_{Fg}$ | Minimum temperature required for egg maturation in females (°C) | 10 | [1] |
| $TDD_{Fg}$ | Total degree-days required for egg maturation (°C) | 77 | [1] |
| $c_L$ | Standard rainfall-independent carrying capacity for larvae $^{-(km^2)}$ | 23629 | [4] |
| $c_P$ | Standard rainfall-independent carrying capacity for pupae $^{-(km^2)}$ | 23629 | [4] |
| $\delta L_a$ | Coefficient a of the larvae mortality function | -0.1305 | [5] |
| $\delta L_b$ | Coefficient b of the larvae mortality function | 3.86 | [5] |
| $\delta L_c$ | Coefficient c of the larvae mortality function | 30.83 | [5] |
| $\delta P_a$ | Coefficient a of the pupae mortality function | -0.1502 | [5] |
| $\delta P_b$ | Coefficient b of the pupae mortality function | 5.057 | [5] |
| $\delta P_c$ | Coefficient c of the pupae mortality function | 3.517 | [5] |
| $T_0$ | Optimum temperature for mosquitoes survival | 22 °C | [6] |
| $p$ | Exponent of the adult-mortality function Hill-like | 6 | Optimization |
| $v$ | Exponent of the adult-mortality function Hill-like | 10 | Optimization |
| $f_E$ | Transition function from egg to larva | Eq. (3) | [2, 7] |
| $m_L$ | Larvae-mortality function | Eq. (10) | [8, 5] |
| $f_L$ | Transition function from larva to pupa | Eq. (4) | [2] |
| $m_P$ | Pupae-mortality function | Eq. (15) | [8, 5] |

Continued on next page

| Notation | Description | Expression | Reference |
|---|---|---|---|
| $f_P$ | Transition function from pupae to emerging adult | Eq. (5) | [2] |
| $m_A$ | Adult mortality | Eq. (??) | Optimization |
| $f_{Fo}$ | Transition function from ovipositing to host-seeking female | Eq. (17) | [2] |
| $f_{Fg}$ | Transition function from engorged to oviposition site-seeking female | Eq. (16) | [2] |
| $f_F$ | Transition rate from nulliparous to parous females | Eq. (18) | [9] |
| $\mu_{Mj}$ | Sterile and toxic males mortality | 0.087 | [7, 10, 3] |
| $c$ | Competitivness of sterile and toxic males | 0.5 | Optimization |

Table 2: Parameters and functions of the SEIR-M model.

| Notation | Description | Expression | Reference |
|---|---|---|---|
| $B_h$ | Function of susceptibility to infections for humans | Eq. (20) | [11] |
| $B_v$ | Function of susceptibility to infections for mosquitoes | Eq. (20) | [11] |
| $\epsilon$ | Human extrinsic incubation period (day$^{-1}$) | 0.25 | Optimization |
| $d_H$ | Human mortality rate given by CHIKV (day$^{-1}$) | $10^{-4}$ | [12] |
| $\rho$ | Human recovery rate (day$^{-1}$) | 0.12 | [13, 14, 15] |
| $\omega$ | Human host preference | 0.6 | [16, 17, 18] |
| $V$ | Biting rate (day$^{-1}$) | Eq. (21) | [11] |
| $\chi_{Bh}$ | Constant rate for Brière function | 4 x $10^{-4}$ | Optimization |
| $\chi_{Bv}$ | Constant rate for Brière function | 5 x $10^{-4}$ | Optimization |
| $\chi_V$ | Constant rate for Brière function | 4 x $10^{-4}$ | Optimization |

###### 1.3.2 Functions

Transition function from egg to larva:

$$f_E = \frac{T(t) - T_E}{TDD_E}, \quad if \quad T(t) > T_E, \quad otherwise = 0. \tag{3}$$

Transition function from larva to pupa:

$$f_L = -0.0007T^2 + 0.0392T - 0.3911. \tag{4}$$

Transition function from pupa to emerging adult:

$$f_P = -0.0008T^2 - 0.0051T + 0.0319. \tag{5}$$

Larvae mortality (Figure 1):

$$T_{\text{opt}} = \frac{\delta L_b}{2|\delta L_a|}, \tag{6}$$

$$T_{\text{eff}} = T_{\text{opt}} + 1.35\,(T - T_{\text{opt}}), \tag{7}$$

$$L = -|\delta L_a|\,T_{\text{eff}}^2 + \delta L_b\,T_{\text{eff}} + \delta L_c, \tag{8}$$

$$L_{\text{eff}}(T) = 16 + \frac{1}{2}\ln\Big(1 + e^{2L(T)}\Big), \tag{9}$$

$$m_L(T) = \frac{1}{L_{\text{eff}}(T)}. \tag{10}$$

Pupae mortality (Figure 1):

$$T_{\text{opt}} = \frac{\delta P_b}{2|\delta P_a|}, \tag{11}$$

$$T_{\text{eff}} = T_{\text{opt}} + 1.05\,(T - T_{\text{opt}}), \tag{12}$$

$$P = -|\delta P_a|\,T_{\text{eff}}^2 + \delta P_b\,T_{\text{eff}} + \delta P_c, \tag{13}$$

$$P_{\text{eff}}(T) = 3.5 + \frac{1}{2}\ln\Big(1 + e^{2P(T)}\Big), \tag{14}$$

$$m_P(T) = \frac{1}{P_{\text{eff}}(T)}. \tag{15}$$

Transition function from engorged to oviposition site-seeking female:

$$f_{Fg} = \frac{T(t) - T_{Fg}}{TDD_{Fg}}, \quad if \quad T(t) > T_Fg, \quad otherwise = 0. \tag{16}$$

Transition function from ovipositing to host-seeking female:

$$f_{Fo} = \gamma_{Fo}(1 + P_{norm}). \tag{17}$$

Transition rate from nulliparous to parous females:

$$f_F = \frac{1}{\frac{1}{f_{Fo}} + \frac{1}{\gamma_{Fh}} + \frac{1}{f_{Fg}}} \tag{18}$$

Adult mortality Figure 2:

$$m_A(T) = \frac{|T - T_0|^p}{|T - T_0|^p + v^p} \tag{19}$$

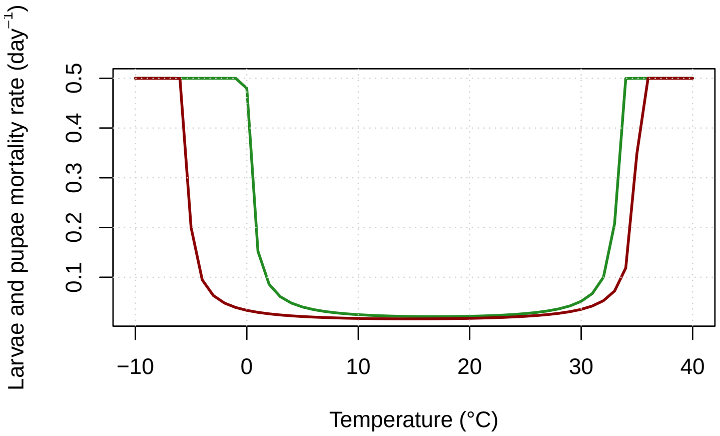


Figure 1: Larvae and pupae mortality functions, described in Eq. (10) and Eq. (15) respectively. In red the curve of mortality of larvae, in green the pupae one.

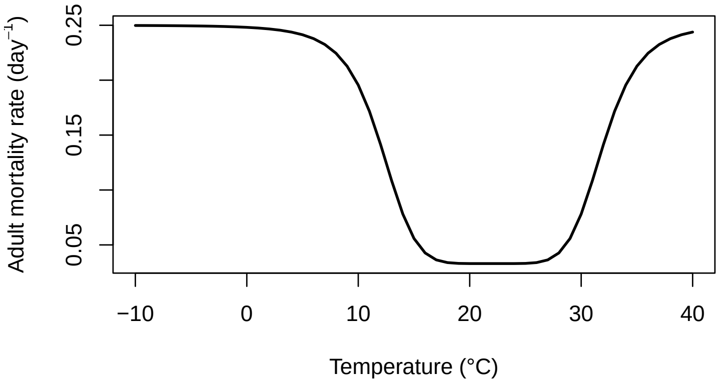


Figure 2: Mortality function of the adult individuals of *Ae. albopictus*, described in Eq. (19).

Brière function ($B_h$ and $B_v$) (Figure 3):

$$B_j = \chi_j T(T - 13.35)(\sqrt{40.08 - T}). \tag{20}$$

Brière function ($V$) (Figure 3):

$$V = \chi_j T(T - 10)(\sqrt{41 - T}). \tag{21}$$

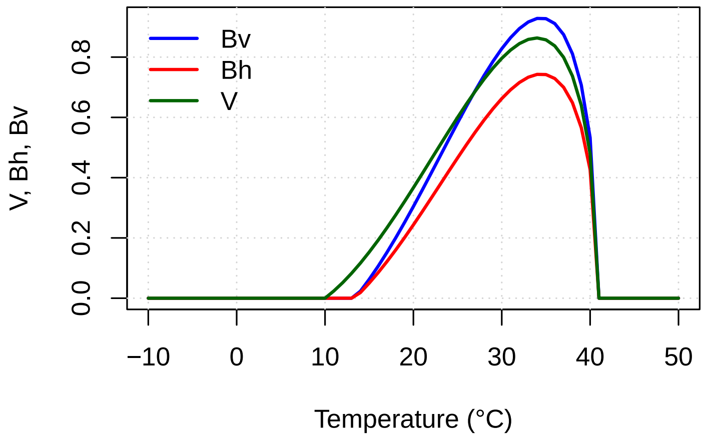


Figure 3: Brière functions defined in Eq. (20). In blue the transmission probability of the pathogen from human to mosquito, in red from mosquito to human, and green line is the biting rate.

Average number of infected humans by a single infective mosquito:

$$R_{vh} = V\omega \frac{Bh}{m_A}. \tag{22}$$

Average number of infected mosquitoes by a single infective human:

$$R_{hv} = V\omega \frac{Bv}{\rho} \frac{\epsilon}{\epsilon + m_A} \frac{N_V}{N_H}, \tag{23}$$

where $N_V$ are the total females mosquito for each time $t$;

Mosquito extrinsic incubation period of the virus:

$$\eta = 1.03(4 + exp(5.15 - 0.123T)). \tag{24}$$

#### 2 Validation of the model

##### 2.1 Emilia-Romagna

To validate our model, we compared the VecAbundance dataset [19] with the weekly cumulative sum of eggs for each time $t$, using the equation:

$$E(t_7) = f_F(\beta_1 F_n(t_7) + \beta_2 F_p(t_7)). \tag{25}$$

The correlation between the two variables was quantified with Pearson coefficient (Figure 4).

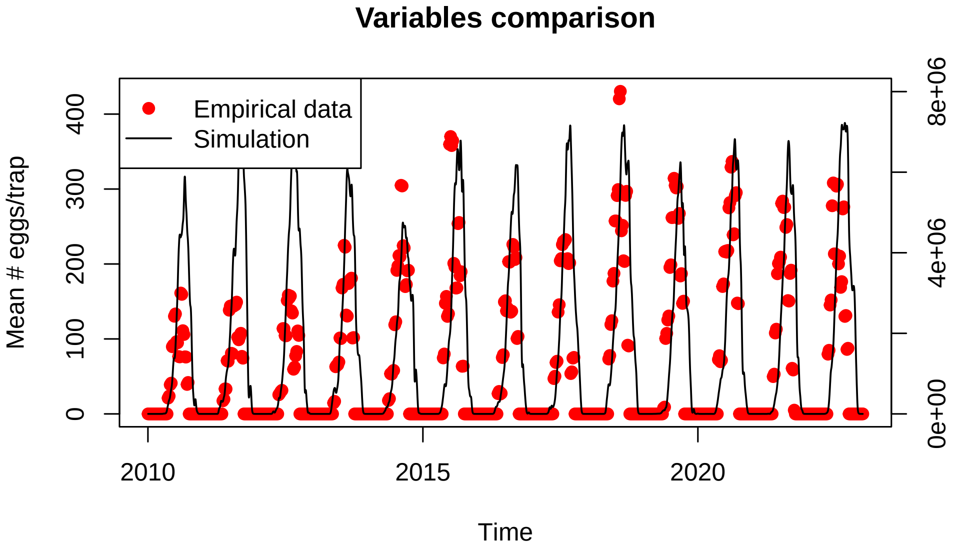


Figure 4: Model validation with cumulative sum of eggs of Emilia-Romagna, Italy. Red dots are observed data from the VecAbundance dataset (left y-axis), and the black line represents the output of the model (right y-axis).

From Figure 4, the predicted egg abundance is consistent with the dataset, with Pearson correlation coefficient r=0.88, p<$10^{-6}$).

##### 2.2 Trentino-Alto Adige

To further demonstrate the reliability of our model, we validated it using Trentino-Alto Adige meteorological data, compared with the weekly cumulative sum of eggs of that region, always extracted from the VecAbundance dataset.

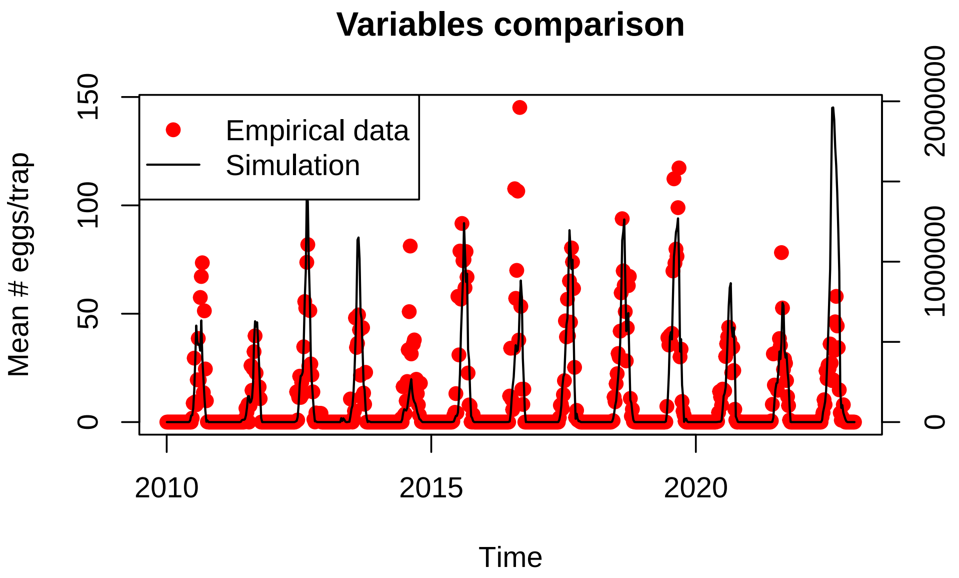


Figure 5: Model validation with cumulative sum of eggs of Trentino-Alto Adige, Italy. Red dots are observed data from the VecAbundance dataset (left y-axis), and the black line represents the output of the model (right y-axis).

As for the Emilia-Romagna case, our simulations align with the data of Trentino-Alto Adige (Figure 5), with a Pearson correlation coefficient r=0.77, p<$10^{-6}$.

[3] Clelia F Oliva, Marco J Maier, Jeremie Gilles, Maxime Jacquet, Guy

Lemperiere, Serge Quilici, Marc JB Vreysen, François Schooneman, Dave D Chadee, and Sebastien Boyer. Effects of irradiation, presence of females, and sugar supply on the longevity of sterile males aedes albopictus (skuse) under semi-field conditions on reunion island. *Acta tropica*, 125(3):287–293, 2013.

[11] John H Huber, Marissa L Childs, Jamie M Caldwell, and Erin A Mordecai. Seasonal temperature variation influences climate suitability for

dengue, chikungunya, and zika transmission. *PLoS neglected tropical diseases*, 12(5):e0006451, 2018.
